# Reactivation of time cell sequences in the hippocampus

**DOI:** 10.1101/389874

**Authors:** Hindiael A. Belchior, Rodrigo Pavão, Alan M.B. Furtunato, Howard Eichenbaum, Adriano B.L. Tort

## Abstract

The temporal order of an experience is a fundamental property of episodic memories, yet the mechanism for the consolidation of temporal sequences in long-term memory is still unknown. A potential mechanism for memory consolidation depends on the reactivation of neuronal sequences in the hippocampus. Despite abundant evidence of sequence reactivation in the formation of spatial memory, the reactivation of hippocampal neuronal sequences carrying non-spatial information has been much less explored. In this work, we recorded the activity of time cell sequences while rats performed multiple 15-s treadmill runnings during the intertrial intervals of a spatial alternation memory task. We observed forward and reverse reactivations of time cell sequences often occurring during sharp-wave ripple events following reward consumption. Surprisingly, the reactivation events specifically engaged cells coding temporal information. The reactivation of time cell sequences may thus reflect the organization of temporal order required for episodic memory formation.

---

Spatially and temporally organized behaviors are accompanied by sequences of neuronal activity in the rat hippocampus (Buzsáki & Moser, 2013; Eichenbaum, 2014). While sequences of place cells encode self-location during spatial trajectories through the environment (O’Keefe & Dostrovsky, 1971; Wilson & McNaughton, 1993), time cell sequences encode successive periods in temporally structured experiences (Kraus, Robinson, White, Eichenbaum, & Hasselmo, 2013; MacDonald, Lepage, Eden, & Eichenbaum, 2011; Pastalkova, Itskov, Amarasingham, & Buzsaki, 2008). Together, the activity of place cells and time cells may establish the spatiotemporal structure necessary for episodic memory representation in the hippocampus (Buzsáki & Moser, 2013; Eichenbaum, 2014). Yet, the neural mechanisms that support the consolidation of spatial and temporal experiences remain only partially understood.

One possible mechanism that promotes the consolidation of hippocampal-dependent memories relies on offline reactivations of experience-related neuronal sequences (see Carr, Jadhav, & Frank, 2011 for a review). Many studies have reported that place cells representing the spatial trajectories previously executed by the animals are spontaneously reactivated during subsequent consummatory behaviors, immobility and sleep episodes (Diba & Buzsáki, 2007; Foster & Wilson, 2006). The reactivation of place cells may occur in forward or reverse sequential order as during spatial navigation, but in a temporally compressed manner through short bursts of neuronal activity frequently associated to hippocampal sharp-wave ripples (100-250 Hz) (Buzsáki, 1998). Although sequence reactivation has been proposed as a general mechanism for memory consolidation, the reactivation of neuronal sequences coding the temporal order of an experience remains much less characterized (but see Wang, Roth, & Pastalkova, 2016).

In this work, we used 24 independently movable tetrodes to bilaterally record neuronal ensembles from dorsal CA1 in five male Long-Evans rats (Supporting Information Fig. S1, see Detailed Methods in Supporting Information). The animals performed a spatial alternation memory task in an 8-shape maze coupled to a computer-controlledtreadmillplacedinthestem(Figure 1A). The task required water-deprived animals to run on the treadmill for 15 seconds on each intertrial interval to obtain water reward at the end of the arms and at the beginning and end of treadmill runnings. The animals were pre-trained on the task to achieve at least 40 correct spatial alternation laps in one session before microdrive implantation surgery. After animals fully recovered from surgery, tetrodes were lowered and positioned close to CA1 pyramidal layer based on electrophysiological parameters, i.e., sharp-wave ripples in the local field potentials. A total of 1159 cells were sorted according to spike waveform features, from which 1125 were classified as putative pyramidal cells and 34 as interneurons based on their mean firing rates and confirmed by their autocorrelograms (Supporting Information Fig. S2). To verify whether pyramidal cells showed temporal specificity during treadmill running, we computed Shannon’s mutual information for each cell as described in Souza, Pavão, Belchior, & Tort (2018).

**Figure 1.**
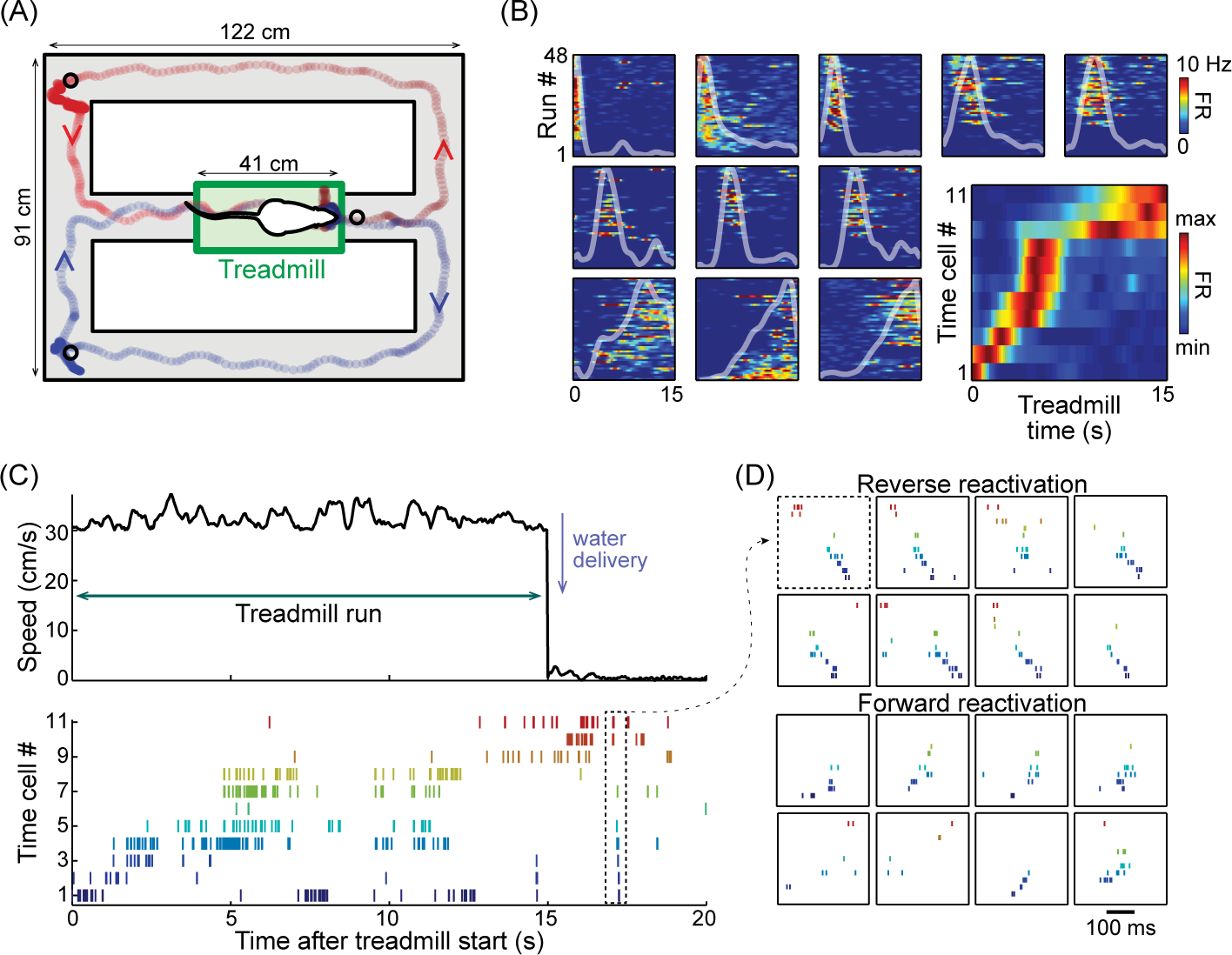
Hippocampal time cell sequences. (A) Task design. Rats ran at 30 cm/s on a treadmill (green rectangle) during a 15-s delay of a spatial alternation task in an 8-shaped maze. Red and blue lines indicate left and right trajectories, respectively. Black circles indicate the three sites of water reward. (B) Color plots show the activity of individual hippocampal time cells across 48 treadmill runs in one experimental session; white lines show the average firing rates. The bottom right panel depicts the template sequence; i.e, the normalized firing rates of all 11 time cells, sorted by the latency to peak firing rate during treadmill running. (C) Instantaneous speed (top) and time cell spike rastergram (bottom) during a treadmill run and subsequent water consumption. Notice sequential activity of time cells during treadmill running (same cells as in B). Dashed rectangle highlights a sequence reactivation. (D) Examples of reverse (top) and forward (bottom) reactivations of time cell sequences (same ordering as in B and C).

Consistent with previous reports (Kraus, Robinson, White, Eichenbaum, & Hasselmo, 2013; MacDonald, Lepage, Eden, & Eichenbaum, 2011; Pastalkova, Itskov, Amarasingham, & Buzsaki, 2008), we observed 491 pyramidal cells coding temporal information during treadmill running, a mean of 12.6 time cells per recording session. Figure 1B shows the spiking activity of 11 time cells across 48 treadmill running periods from one experimental session(see also Supporting Information Fig. S3). Only a few cells encoded time since the first treadmill runnings, while most of them established temporal fields along the session, in resemblance to reports about the dynamics of place field formation (Bittner, Milstein, Grienberger, Romani, & Magee, 2017). To construct time cell template sequences, for each recording session we used only time cells exhibiting unimodal time fields. Time cells in each template sequence were then ordered by the latency to peak firing rate within the 15-s treadmill running time. A typical template sequence with ordered time cells is shown in Figure 1B (bottom right panel). Individual time cells showed no difference in firing rate during treadmill running preceding left and right laps on the maze, and template sequences usually covered most of the treadmill running time (Supporting Information Fig. S3 and S4).

In order to identify reactivation candidates, we selected epochs where the treadmill was off and the speed of locomotion of the rat was <1 cm/s. Within these epochs, we analyzed bursts of population spiking activity in which at least 3 cells were member of the time cell template sequence. Only task sessions containing more than 15 reactivation candidates were selected for further analysis. Using these criteria, we found a total of 621 reactivation candidates in ten recording sessions from three rats.

We then examined whether the time cell template sequences observed during treadmill runnings correlated to the order of activated neurons within the bursts of spontaneous activity during quiet behavior. To do that, we calculated the Spearman rank-order correlation coefficient (rho) between spike times observed in each reactivation candidate and the order of time cells in the template sequence (Diba & Buzsáki, 2007). Candidates showing correlation coefficients lower than 2.5% or higher than 97.5% of 500 rho values obtained from shuffled distributions were classified as significant reactivations (Supporting Information Fig. S5).

Figure 1C shows the spiking activity of an ensemble of time cells during a typical treadmill running and the following consummatory behavior. Note that the sequential activation of time cells during treadmill running reflects the template sequence shown in Figure 1B (same cell order on Y-axis). The example also shows a burst of spontaneous spiking activity in the same population of neurons just after the treadmill stopped, which was classified as a reactivation candidate and further analyzed. Additional analysis revealed that this example was a significant reverse reactivation of the time cell template sequence. Other examples of significant reverse and forward reactivations of time cell sequences are shown respectively in the top and bottom panels of Figure 1D (see also Supporting Information Fig. S5). Similar to previous reports (Lee & Wilson, 2002; Wang, Roth, & Pastalkova, 2016), the reactivation of time cell sequences were temporally compressed, lasting around 200 ms (see Supporting Information Fig. S6). These results show that sequences of time cells are reactivated during bursts of spiking activity in both forward and reverse directions.

Group analysis of ten sessions from three rats showed 93 reverse and 81 forward reactivations (15% and 13%, respectively). Figure 2A shows the distribution of significant correlations estimated using the standard spike-shuffling procedure originally employed to detect reactivations of place cell sequences (Diba & Buzsáki, 2007; Dragoi & Tonegawa, 2011, 2013; Silva, Feng, & Foster, 2015). We observed that 28% of the reactivation candidates were deemed significant under this analysis, thus exceeding the 5% expected by chance. To further investigate the robustness of reactivation events, we performed an additional statistical testing which consisted of shuffling neuronal identities in the template sequence (Supporting Information Fig. S5). In this procedure, we kept the reactivation candidates in the original order, while checking for correlations with templates in which the order of neurons was shuffled. If this procedure were unbiased, the percentage of candidates correlated with the shuffledtemplatewouldbeatthe 5% chancelevel. However, we found that shuffled template sequences displayed more reactivations than expected by chance (Figure 2B), indicating that the standard spike-shuffling procedure is biased towards false positives and that previous studies were likely to have overestimated reactivation rate. Importantly, the incidence of reactivation observed with the actual template was significantly higher than with the shuffled template sequences (Figure 2B), confirming that time cell sequences reactivate at a greater extent than chance.

**Figure 2.**
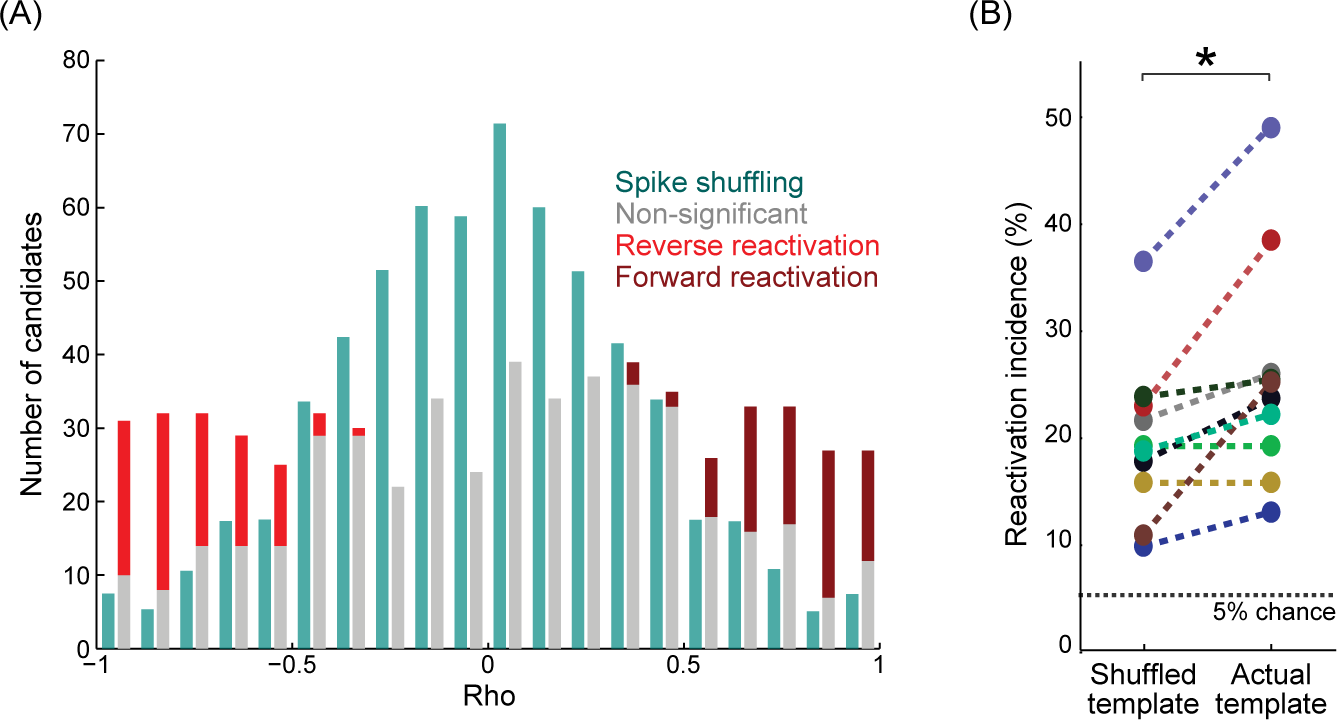
Time cells exhibit reverse and forward sequence reactivation. (A) Rank-order correlation coefficients between reactivation candidates and template sequences (n=621 candidates across 10 sessions). Candidates were classified as reverse (15%, light red) or forward (13%, dark red) reactivations based on standard spike-shuffling methods (Diba & Buzsáki, 2007; Dragoi & Tonegawa, 2011, 2013; Silva, Feng, & Foster, 2015). Green bars show the normalized distribution of correlation coefficients for shuffled spikes. (B) Reactivation incidence obtained through spike-shuffling method when using shuffled template sequences (left) or actual templates (right) in the correlation analysis (see Supporting Information Fig. S5). Notice that the reactivation incidence found for shuffled templates is greater than 5% for all sessions. Nevertheless, reactivation incidence was higher for the actual template (P=0.0078, Wilcoxon signed-rank test). Colors denote different sessions.

To further characterize the reactivation of time cell sequences, we investigated the position of the rat on the maze and the time in which reactivation events occurred. We found that reactivations occurred close to the water ports, with no spatial difference between reverse and forward reactivations (Figure 3A). Most reactivation events followed reward consumption, about 5 s after the treadmill stopped or the animals reached the reward sites in the arms (~30 s afterward) (Figure 3B). We also observed anincrease in the rate of reactivationevents along each recording session (Figure 3C), which may reflect the establishment of time fields along the task, as described above; alternatively, this may indicate the strengthening of the sequences of time cells reactivated during repetitive and rewarded treadmill runnings (Singer & Frank, 2009).

**Figure 3.**
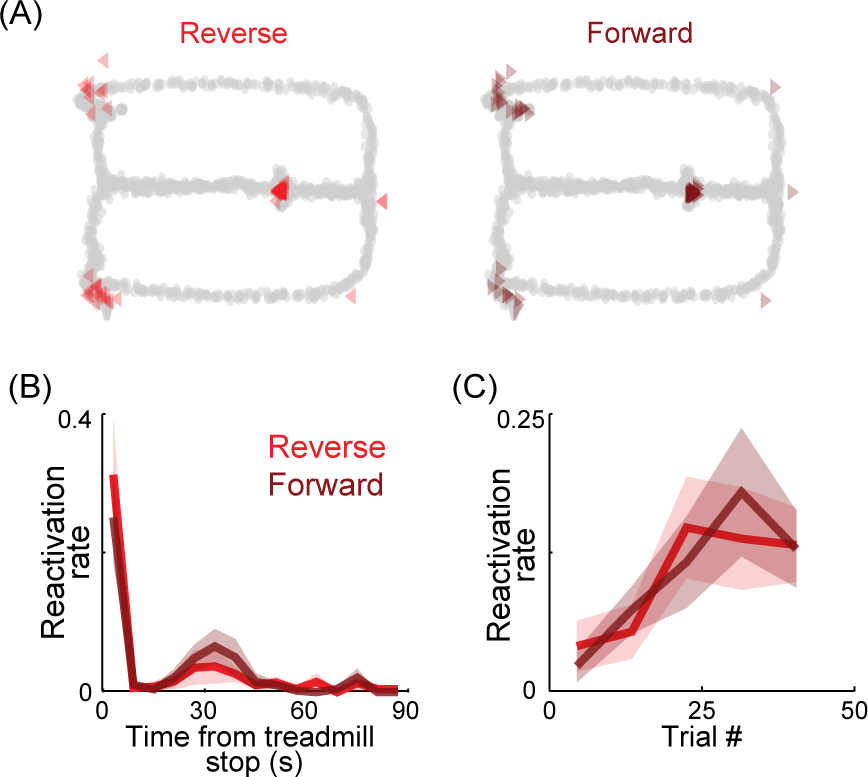
Temporal and spatial features of time cell sequence reactivations. (A) Spatial distribution of reverse (left) and forward (right) reactivation events for all the 10 recording sessions. Notice that reactivations occur at reward locations in central, left and right reward locations. (B) Normalized reactivation rate as a function of time from treadmill stop. Both reverse and forward reactivations peak immediately after treadmill stop and ~30 s afterward, during water consumption. (C) Normalized reactivation rate as a function of trial number. To allow merging across different sessions, the number of reactivations in a trial bin was divided by the total number of reactivations of the session. Results show mean ± S.D. over 10 sessions. Notice that reactivation rate increases within sessions.

We then investigated how time cells and other cell types convey temporal and spatial information. Figure 4A shows the amount of temporal information (time on treadmill) and spatial information (position on the maze) conveyed by firing rates of time cells (left panel), not-time cells (middle panel) and interneurons (right panel). We found that 97.5% of time cells (76 units) also carried spatial information along the maze, and that the amounts of temporal and spatial information were not correlated. We observed that 58.2% of the “not-time cells” (110 units) conveyed spatial information. Interestingly, all interneurons (16 units) conveyed significant and correlated amount of temporal and spatial information. These results confirm that time cells may encode non-temporal aspects of the experience (Kraus, Robinson, White, Eichenbaum, & Hasselmo, 2013), and show that interneurons encode both spatial and temporal information. We then investigated how these cell-types engage in the reactivation events. As expected, time cells increased their firing rate at reactivation events (Figure 4B). Conversely, the firing rates of “not-time cells” were not significantly modulated during the reactivation of time cell sequences. Surprisingly, interneurons also increased their firing rate during reactivations events (Figure 4B). Finally, in agreement with reports of place cell sequence reactivation (Diba & Buzsáki, 2007; Foster & Wilson, 2006), time cell sequences also reactivated accompanied by sharp-wave ripple events in the local field potential (Figure 4C and Supporting Information Fig. S6). These findings are consistent with the hypothesis that time cells and place cells share similar reactivation mechanisms associated with sharp-wave ripples (Buzsáki, 2015), but also indicate that temporal and other aspects of an experience may be encoded by distinct reactivation events.

**Figure 4.**
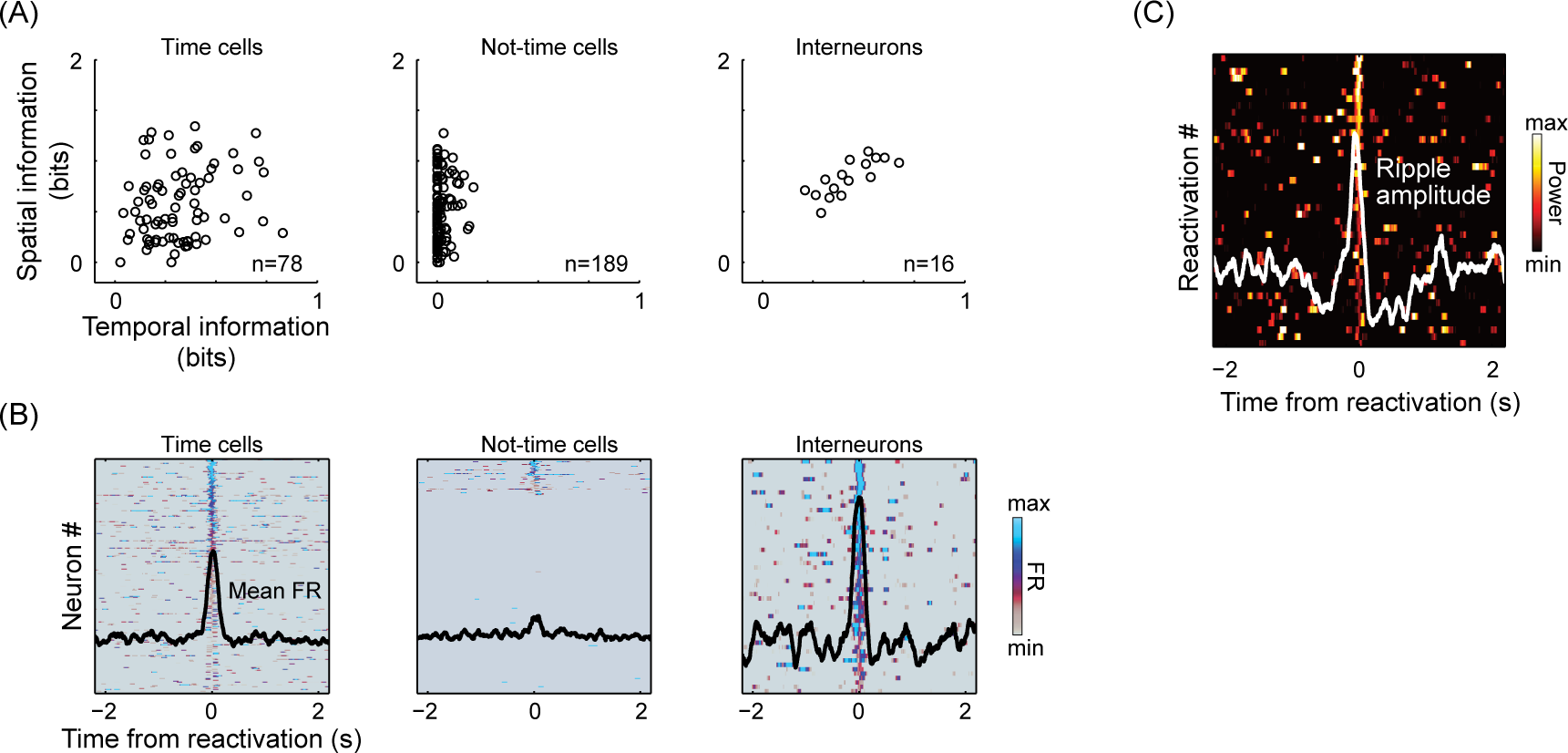
Cell-type specificity and ripple activity and during time cell sequence reactivation. (A) Circles depict temporal and spatial information conveyed by the firing rate of each cell. (Left) Time cells encoded temporal information on the treadmill but also spatial information on the maze. Notice the amount of information encoded by the time cells on the treadmill did not correlate with the information encoded on the maze (rho=0.21). (Middle) “Not-time cells”, that by definition do not encode temporal information, encode varying amount of spatial information on the maze. (Right) Interneurons encoded both temporal and spatial information, in related amounts (rho=0.78). (B) Firing rates of time cells, “not-time cells” and interneurons around reactivation events. Each row represents the normalized firing rate of one neuron on each reactivation. Only the highest firing rates at reactivations are shown (top 20%). Black lines depict mean firing rate. Notice that only time cells and interneurons increase firing during sequence reactivations. (C) Normalized ripple amplitude around time cell sequence reactivations (top 20% events); white line shows mean ripple amplitude.

Our results provide evidence for the forward and reverse reactivation of time cell sequences in the hippocampus. The reactivation of time cell sequences occurred during quiet and consummatory behaviors and associated with sharp-wave ripples. This finding parallels previous reports of sequence reactivation in hippocampal neurons representing spatial trajectories, b ut for the temporal domain (Diba & Buzsáki, 2007; Foster & Wilson, 2006). Although a recent report has shown that the formation of place and time cell sequences differs (Wang, Roth, & Pastalkova, 2016), the reactivation of temporal and spatial sequence during sharp-wave ripples depends on similar mechanisms. In addition, our results suggest that the occurrence of reactivations containing only time cells supports the independent processing of temporal experiences in blocks of selective reactivation events (Gupta, van der Meer, Touretzky, & Redish, 2012).

In summary, here we report that hippocampal time cell sequences reactivate in reverse and forward order during sharp-wave ripples. Along with spatial sequences, reactivation of temporal sequences may strengthen interconnectedneuronalensemblescodingspatiotemporal features of episodic memories. These results indicate that temporal and spatial neuronal sequences may rely on the same neural mechanisms to consolidate long-term hippocampal-dependent memories (Buzsáki & Moser, 2013; Eichenbaum, 2014).

## Methods

### Animal experimentation

This study was performed in strict accordance with the recommendations in the Guide for Care and Use of Laboratory Animals of the National Institutes of Health. All experimental protocols were approved by the Boston University Institutional Animal Care and Use Committee.

### Behavioral protocol

We used a modified eight-shape maze (122 cm x 91 cm) in which the central stem contained a computer-controlled treadmill (41 cm x 12 cm, Columbus Instruments). Animals were trained to execute a spatial alternation memory task associated to treadmill running during intertrial intervals for a period of 15 s at constant speed of 30 cm/s. Animals were water deprived for 12 h before experiments. Water drops (0.05 ml) were delivered in water ports located ahead of the treadmill and at the end of each alternation arm (Figure 1A). Before surgery for electrode implantation, each animal performed at least 35 successful alternation laps during 3 consecutive training sessions. Experimental sessions were recorded through a digital video camera (30 frames per second) placed above the maze and synchronized to electrophysiological recordings.

### Surgery and electrophysiological recordings

Following training, five male Long-Evans rats (400-550g) were bilaterally implanted with 24 independently movable tetrodes targeting the dorsal CA1 area of the hippocampus (3.2 mm AP and 1.9 mm ML relative to Bregma, Supporting Information Fig. S1) under 1-3% isoflurane anesthesia. Tetrodes consisted of four twisted 12.7 µm polyimide-insulated nickel-chromium wires (Sandvik) electroplated in gold solution to reduce impedance to ~200 kOhms at 1 kHz using NanoZ (Neuralynx). At the end of surgery, tetrodes were lowered ~1 mm into brain tissue. After recovery, tetrodes were progressively lowered to the pyramidal cell layer during two weeks. Final tetrode positioning was estimated based on stereotaxic coordinates and electrophysiological benchmarks. One tetrode in each hemisphere was placed at the external capsule of the corpus callosum and used as reference for ipsilateral tetrodes. One tetrode was further lowered and positioned close to the hippocampal fissure to record high-amplitude theta oscillations. All other tetrodes were positioned close to pyramidal cell layer, as indicated by the occurrence of ripple events, complex-spikes and theta-modulated spiking activity. Electrodes were never moved within the 12 hours preceding recording sessions. Electrophysiological recordings were performed using a multichannel acquisition processor (96-channel MAP System, Plexon Inc.). Brain signals were amplified (1,000-10,000 x), filtered for high-frequency spiking activity (154-8.8 kHz) and for local field potentials (1.5-400 Hz). Local field potentials were digitized at 1 kHz; spiking activity was detected by threshold crossing and waveforms (0.8 ms) were digitized at 40 kHz. Two light-emitting diodes located at the headstages were used to track animal position.

### Histology

After experiments, rats were overdosed with pentobarbital (100 mg/Kg, i.p.), and an electrical current (25 µA) was applied for 30 s to mark brain tissue around the tetrode tips. Animals were then perfused with 0.9% NaCl solution, followed by Prussian blue solution (10% potassium ferrocyanide and 4% paraformaldehyde solution in phosphate buffer 0.1 M, pH of 7.4). Brains were removed, maintained in 20% sucrose solution for 24 h before frozen. Coronal sections (50 µm) were performed in a cryostat (Micron) and mounted over glass slides. Brain sections were stained with cresyl-violet solution. Electrode location was estimated by light microscopy (Supporting Information Fig. S1B).

**Figure S1.**
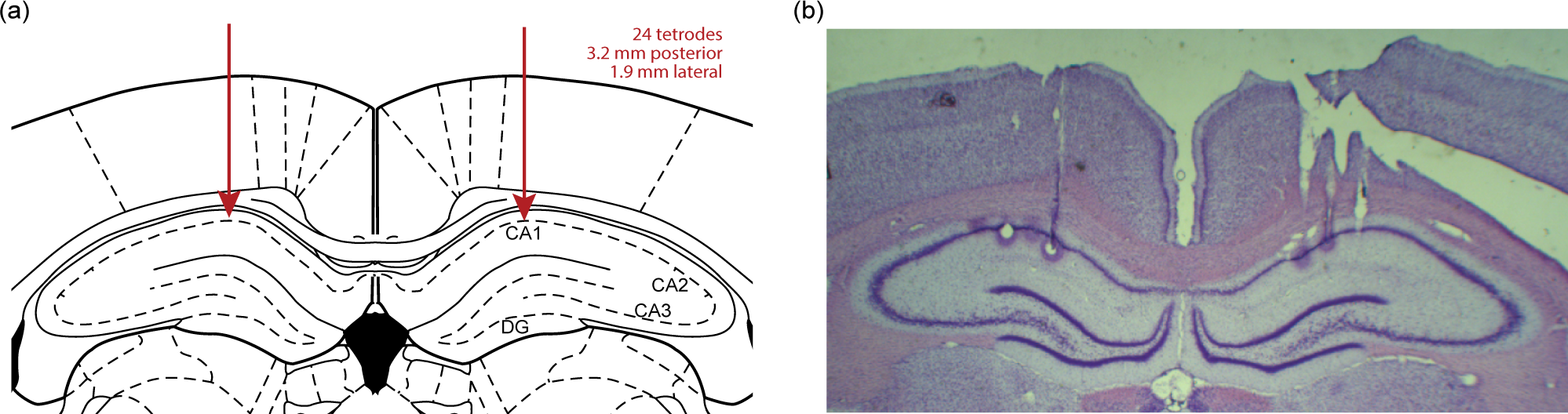
Stereotaxic coordinates and representative histology. (A) Schematic illustration of a coronal section of the rat brain at 3.2 mm posterior from Bregma. Red arrows indicate hippocampal coordinates used as recording targets (1.9 mm lateral from the midline). (B) Representative cresyl-stained coronal brain section. Electrical current lesions indicate the location of the tetrode tips at the CA1 pyramidal cell layer.

### Data analysis

#### Spike sorting

Individual neuronal units were manually isolated using Offline Sorter (Plexon Inc.). For each tetrode, clustering was based on differences in spike waveform features (i.e., peak, amplitude, and peak-valley difference) across recording channels (Supporting Information Fig. S2A and S2B).

Putative pyramidal cell and interneuron classification. Single units were classified as putative pyramidal cells and interneurons based on mean firing rates. Neurons with firing rates higher than 8 Hz were classified as interneurons. We then computed the autocorrelogram of each cell to confirm whether putative pyramidal cells exhibited spikes in bursts and putative interneurons exhibited regular spiking activity (Supporting Information Fig. S2C and S2D).

#### Temporal and spatial information

Continuous firing rate time series (dt=1 ms) were obtained by convoluting spike trains of pyramidal cells with a gaussian kernel (σ=450 ms). During treadmill running, firing rate values were binned into 10 equi-populated bins; treadmill time was binned into non-overlapping 150-ms bins. Neurons were classified as time cells when the Shannon’s mutual information between their firing rate and treadmill time was higher than chance (>99% of 1000 surrogate information values computed using shuffled time bins) (Souza, Pavão, Belchior, & Tort, 2018). A similar procedure was used to quantify spatial information, in which mutual information was calculated between bins of firing rate and spatial location (3.35 cm width) along the maze. Pearson correlation was used to quantify dependence between temporal and spatial information in different cell types (Figure 4B).

**Figure S2.**
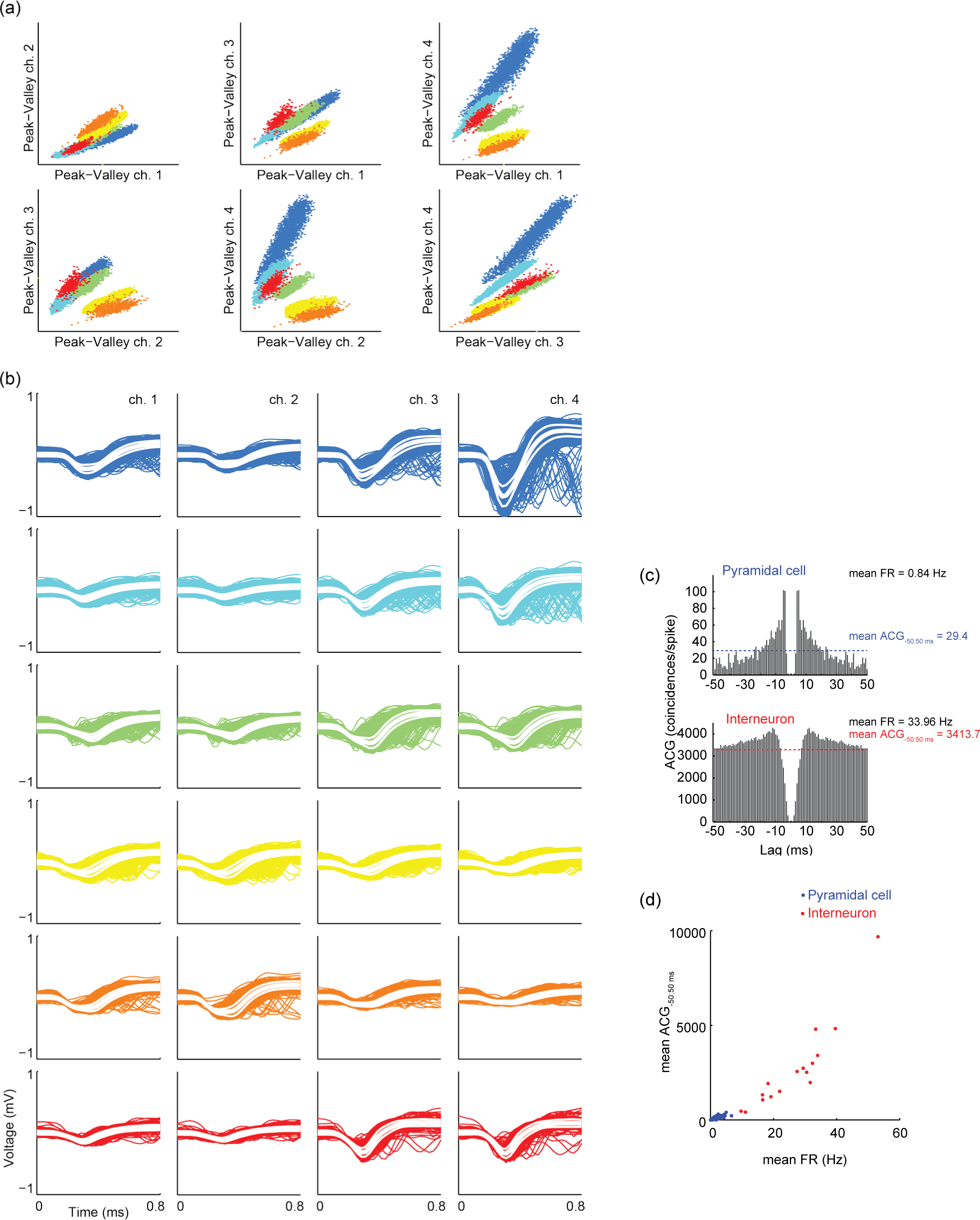
Spike sorting based on waveform features. (A) Representative scatter plots showing peak-valley amplitudes of spike waveforms for all pairwise combinations of tetrode channels; 6 single units were detected (color coded). (B) Waveforms for the same units as in a; white lines indicate mean ± S.E.M. (C) Autocorrelograms for a putative pyramidal cell (top) and a putative interneuron (bottom). Dashed lines indicate the mean autocorrelogram values for each cell. (D) Mean autocorrelogram values as a function of mean firing rates. Notice the separation between the clusters of putative pyramidal cells (blue) and putative interneurons (red).

**Figure S3.**
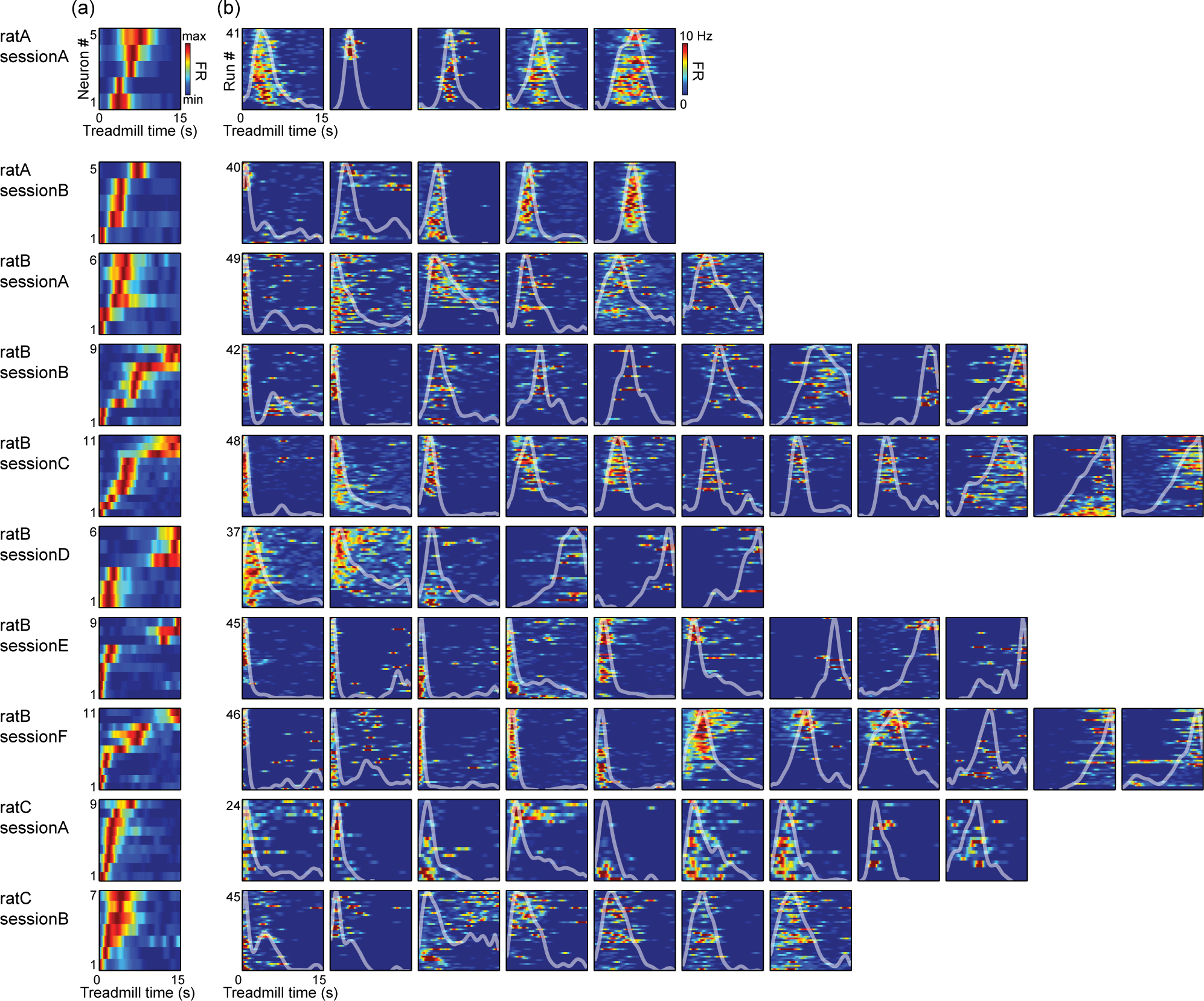
Time cell ensemble activity in each experimental session. (A) Template sequences constructed by normalizing firing rate and ordering time cells along the y-axis by the latency to peak firing rate during treadmill running; the x-axis depicts treadmill running time. Warm colors indicate increased firing rate (FR). Different rows show different sessions, as labeled. (B) Activity of individual hippocampal time cells. For each color plot, the y-axis depicts different treadmill runs. The white lines show the average firing rate as a function of treadmill time.

#### Time cell template sequences

Template sequences were constructed using unimodal time cells whose peak firing rate averaged across trials was >1 Hz (“template time cells”). Unimodality was defined as no additional firing rate peak larger than half the amplitude of the main peak. For each cell, the latency to peak firing rate after the treadmill start was used to order the template sequence (Supporting Information Fig. S3). To test whether time cells had goal-direction selectivity, we compared firing rates during the treadmill run preceding left and right spatial alterations. We used Wilcoxon rank-sum test with significance level of 0.01 (Supporting Information Fig. S4).

#### Reactivation candidates

Reactivation candidates were obtained similarly to previous work (Diba & Buzsáki, 2007; Dragoi & Tonegawa, 2011; 2013; Silva, Feng, & Foster, 2015). Namely, we selected periods with more than five active pyramidal cells, 3 of which template time cells, preceded and followed by 100 ms with no pyramidal cell spikes. We evaluated only periods of quiet wakefulness with locomotion speed lower than 1 cm/s for at least 500 ms (Supporting Information Fig. S5A).

#### Correlation analysis

Only sessions with more than 15 reactivation candidates were included in the correlation analysis. For each reactivation candidate, we calculated the Spearman rank-order correlation coefficient (rho) between spike times and latencies to peak firing rate of the template time cells (Supporting Information Fig. S5B). The rho of each candidate was compared to a surrogate distribution of 500 rho values from shuffled spike identities (Diba & Buzsáki, 2007; Dragoi & Tonegawa, 2011, 2013; Silva, Feng, & Foster, 2015). The candidate was classified as (1) significant reverse reactivation when rho values were lower than the 2.5% of the surrogate distribution; (2) significant forward reactivation when rho values were higher than 97.5%; or (3) non-significant reactivation candidates were between 2.5% and 97.5% of the surrogate distribution (Supporting Information Fig. S5C). The reactivation incidence (RI) corresponds to the percentage of significant reactivations within all candidates (Supporting Information Fig. S5D).

**Figure S4.**
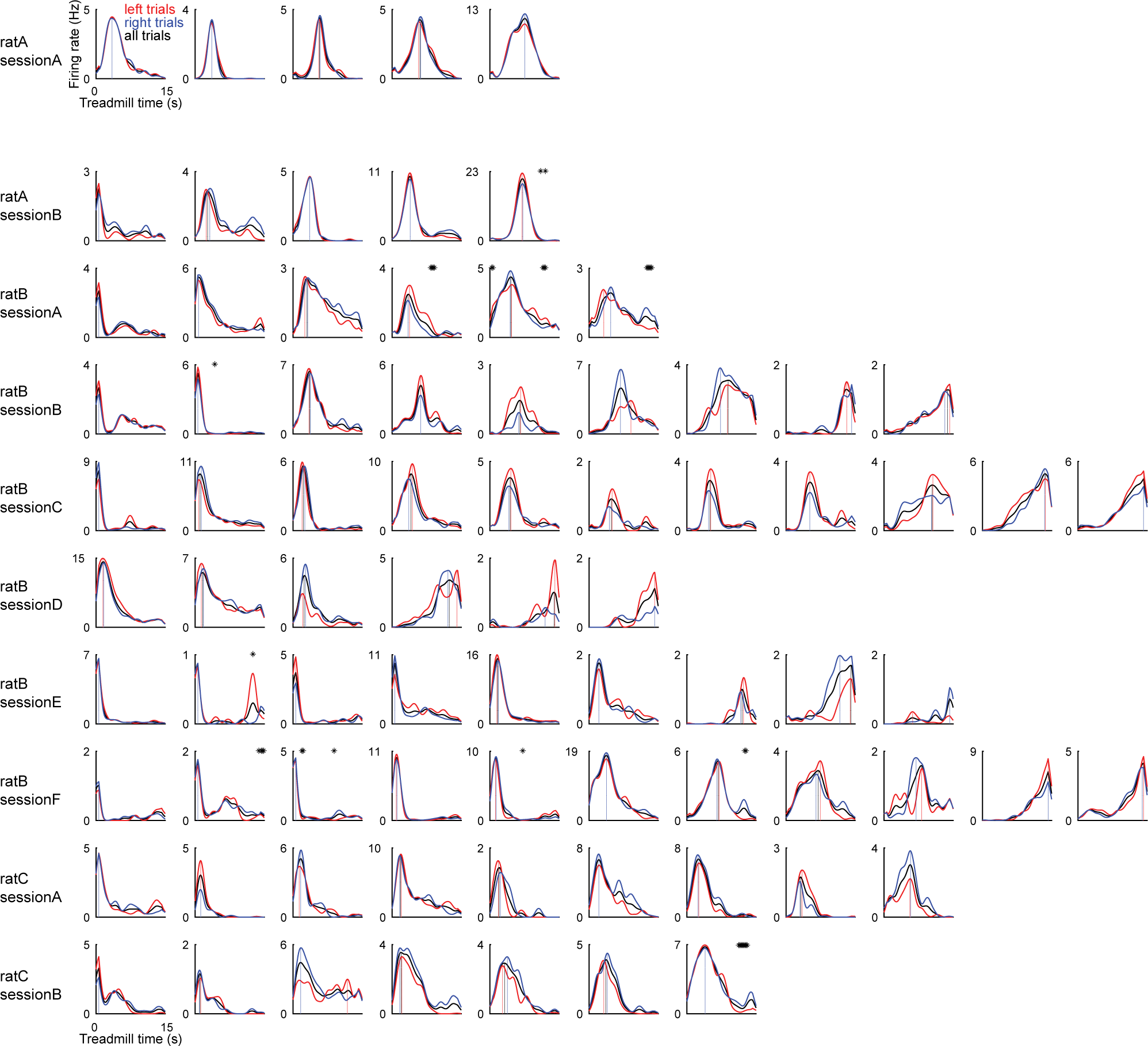
Mean firing rate of template time cells across treadmill runnings. Black lines depict firing rates averaged over all runs, red and blue lines show mean firing rates during treadmill running preceding left and right alternations. Asterisks denote time bins with p<0.01 in Wilcoxon rank-sum test. Notice that most cells do not show different firing rates according to the spatial choice on the following alternation.

#### Template shuffling

The standard spike-shuffling procedure (Diba & Buzsáki, 2007; Dragoi & Tonegawa, 2011, 2013; Silva, Feng, & Foster, 2015) assumes that the RI expected by chance is 5% (the sum of the tails of the surrogate distribution). We tested this assumption by comparing the RI of the actual template sequence with a surrogate RI distribution obtained by 500 shufflings of the identity of time cells in the template sequence. That is, each surrogate RI is computed using the same standard spike-shuffling procedure but applied to a shuffled template sequence. RI distributions above 5% indicate that the spike-shuffling procedure is biased towards false positives (Supporting Information Fig. S5E). We compared actual RI values with the corresponding medians of the template-shuffled distributions using Wilcoxon signed-rank test (Figure 2B).

#### Cell-type specificity

In order to evaluate which cell types (e.g., time cells, not-time cells, interneurons) were modulated during reactivations (Figure 4B), we averaged their z-scored firing rate over all reactivations. Firing rate was computed using 100-ms sliding windows; the z-score normalization was performed within neurons. For each cell type, we compared the distributions of firing rate at the reactivation (t=0 s) and 2 seconds before and afterwards using Kolmogorov-Smirnov test.

#### Ripple amplitude

We band-pass filtered raw local field potential (LFP) signals between 150 and 250 Hz and calculated the instantaneous amplitude using the Hilbert transform. In Figure 4C, we averaged the z-scored ripple amplitude triggered by reactivation events. To that end, for each session we selected the channel with highest ripple amplitude during quiet wakefulness. We compared the distributions of ripple amplitude at the reactivation (t=0 s) and 2 seconds before and afterwards using Kolmogorov-Smirnov test.

**Figure S5.**
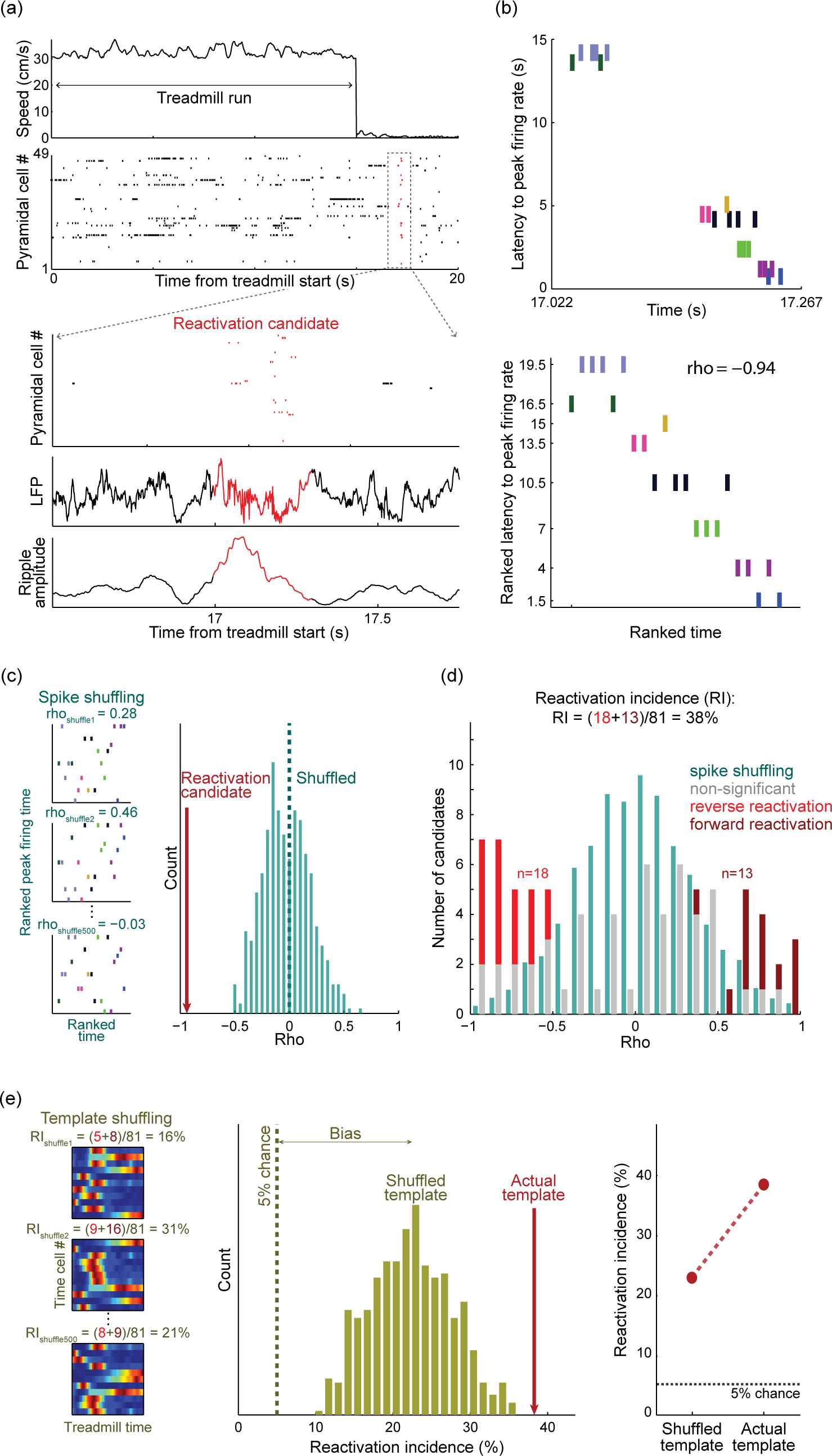
Detecting time cell reactivations. (A) (Top) Instantaneous speed and spike times of putative pyramidal cells recorded during one treadmill run followed by reward consumption. Dashed square highlights spiking activity from one reactivation candidate. (Bottom) The reactivation candidate co-occurred with a ripple event in the LFP, as confirmed by increased amplitude in the ripple frequency band (100-250 Hz) (Figure 4C). (B) Scatter plots show the spike times of the reactivation candidate event in A. For each cell, the y-axis indicates the latency to peak firing rate in the template sequence (see Supporting Information Fig. S3) in parametric (top) and nonparametric (bottom) scales. The rho value refers to the Spearman rank-order correlation coefficient. (C) The left panels show 3 examples of spike shufflings within the reactivation candidate along with corresponding rho values. The right panel shows the distribution of 500 rho values obtained by spike shuffling (cyan), which centers around zero (dashed line). Since the actual rho value (red arrow) is lower than 2.5% of the distribution, the candidate was classified as a reverse reactivation. (D) Histogram of correlation coefficients for all 81 reactivation candidates in the session. Significantly correlated candidates corresponded to 38% (“reactivation incidence”, RI) and are displayed in light red (reverse, n=18) or dark red (forward, n=13); gray denotes non-significant candidates. Cyan bars depict the pool of surrogate distributions obtained by 500 spike shufflings per candidate (rescaled for visual inspection). (E) The left panels show 3 examples of shuffled template sequences, in which neuronal identities in the y-axis were randomly permuted. The middle panel shows the distribution of RI values obtained from shuffled template sequences. Notice that surrogate RI values are greater than 5% (dashed line). The red arrow points to the actual template RI. The right panel shows the average RI for shuffled templates versus the actual template RI. These results were obtained for rat B, session C (same session as in Figure 1).

**Figure S6.**
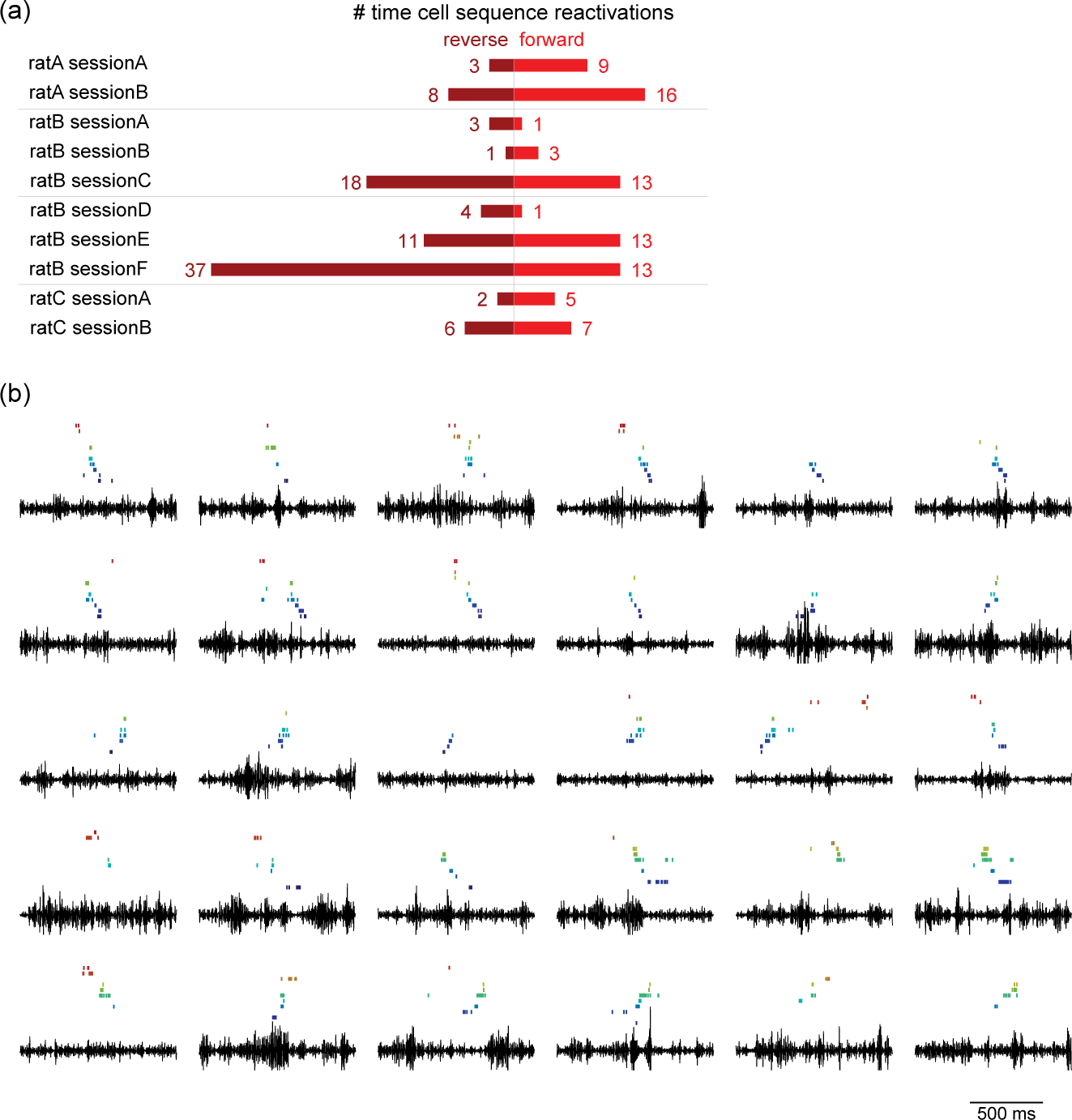
Reverse and forward reactivations of time cell sequences. (A) Number of reverse (dark red) and forward (light red) time cell sequence reactivations per recording session according to standard spike-shuffling criteria. (B) Examples of forward and reverse reactivations and concomitant ripple band (100-250 Hz) activity (black traces). In each panel, cells were ranked by the latency of peak firing rate during treadmill running. Only spikes within reactivations are shown.

## Acknowledgments

This work was supported by the Conselho Nacional de Desenvolvimento Científico e Tecnológico (CNPq), Grant Sponsors: CNPq Universal (N°478331/2013-4), Coordenação de Aperfeiçoamento de Pessoal de Nível Superior (CAPES), and NIMH MH095297. The authors thank Robert J. Robinson II for technical assistance.

